# Computational Assessment of Circularization Impact on Saxitoxin G-quadruplex Aptamer Performance

**DOI:** 10.1101/2025.05.05.652042

**Authors:** Siyi Gao, Bowen Deng, Dingyuan Cheng, Rong Zhou, YuLong Wu, Yuanyu Jia, Boyi Xiao, Haowen Yu, ZhenXia Ma, Yun Gao, TianYin Li, Genkun Liu, Yinuo Zhang, Jun Cheng, Hu Zhu, Han chen, Lianghua Wang, MingJuan Sun

## Abstract

Although high-affinity and high-specificity aptamers can be obtained by SELEX (Systematic Evolution of Ligands by Exponential Enrichment), and have good activity in vitro, their performance in vivo is poor. This is mainly because of the degradation of linear nucleic acids by nucleases in body fluids and the difficulty of maintaining stable 3D structures in complex body fluid environments, which seriously hinders the clinical application of aptamers. This makes circular aptamers, which can resist degradation and are more stable in complex body fluid environments, an attractive option. However, the cyclization process may have a considerable impact on the structure of the original linear aptamer, and blind cyclization may lead to a decrease in its affinity. Here, we developed a new method to guide the cyclization of saxitoxin (STX) aptamers based on molecular computation, and verified the activity of circular aptamers through bioactivity experiments to further verify the authenticity and reliability of our methods. Consistent with the computational analysis, although circularization disrupts the G4 structure, circular aptamer not only has a higher affinity than the linear one but also provides certain protection to mice and greatly prolongs their survival, which proves the accuracy and reliability of the circular structure designed by our computational method. Our findings provide important ideas for better guidance in the design of circular aptamers and demonstrate their great potential in therapeutic applications.

## 1. Introduction

Saxitoxin (STX) is one of the most toxic marine biotoxins and is currently known as one of the main toxins in paralytic shellfish poisoning (PSP). In view of the high hazard and wide distribution of STX, it has been listed as a mandatory item in aquatic product safety inspections worldwide [1]. The lethal dose in humans is approximately 1 mg [2]. Additionally, the mortality rate of STX poisoning is approximately 15% [3]. After 30 min of ingesting food containing STX, a person will feel a tingling sensation on the lips, accompanied by a series of symptoms such as respiratory paralysis, a drop in blood pressure, and even cardiac arrest, which can lead to death within 15 min. No antidote exists for saxitoxin toxicity, and the main clinical treatments are emetic administration and gastric lavage [4].

Although some studies have suggested that 4-aminopyridine can antagonize the toxic effects of STX and provide a therapeutic effect [5], it has highly toxic side effects and a small safe-dose range. Therefore, the therapeutic efficacy and safety of 4-aminopyridine for STX poisoning require further investigation. Due to the lack of a specific antidote, accidental ingestion of seafood containing STX causes many deaths yearly; STX may also have uncertain chronic toxicity [6], which has a huge impact on the marketing of seafood and the normal operations of fishermen. Therefore, it is important to develop an effective and economical STX drug with low toxicity to reduce long-term negative effects and mortality caused by STX poisoning.

Aptamers are oligonucleotide fragments selected from a library of nucleic acids using SELEX technology [7]. They are functional nucleotide molecules with complex three-dimensional (3D) structure. Owing to their unique 3D structure and smaller molecular weights, different aptamers can bind with high specificity and affinity to specific target substances, including some small molecules and ions [8]; thus, aptamers have great potential for both therapeutic and diagnostic applications. Although there are many reports on aptamer-based biosensors with low detection limits and inexpensive preparations [9], pegaptanib (Macugen) is the only nucleic acid aptamer drug marketed for the treatment of age-related macular degeneration [10]. There is still a considerable gap in the therapeutic application of aptamers. Most unmodified aptamers exhibit good activity in vitro, but are susceptible to degradation by nucleases or large conformational changes in the complex in vivo environment, resulting in reduced affinity and targeting ability, which severely hinders the clinical application of aptamers to humans. Recently, chemical modification, such as polyethylene glycosylation (PEGylation), locked nucleic acid (LNA), and 2’-substitutions on sugars (2′-F,2′-NH2 and 2′-O-methyl 2′-OMe), has been reported in many studies to enhance the in vivo stability of aptamers and resist degradation from nucleases [11,12,13,14]. However, such chemical modifications usually require complex organic synthesis. In addition, the chemical modification of aptamers is likely to result in a change in conformation and hinder the binding of the target at the appropriate site, ultimately reducing affinity and pharmacological activity [15,16], Furthermore, chemical modification may also increase the immunogenicity of aptamers [17,18], and some aptamers may become toxic [19,20]; therefore, not all aptamers are suitable for this method. Aptamer cyclization has emerged as a promising alternative for several reasons. First, it maintains the stability of the aptamer and makes it resistant to degradation by ligating the 5′ and 3′ ends [21], similar to chemical modification; second, it circumvents the requirement of complex chemical synthesis simply by using DNA ligase for the ligation process. Third, cyclization preserves the natural state of the aptamer as much as possible, so that the circularized aptamer does not increase toxicity or immunogenicity. Lei et al. screened a circular aptamer, CTBA4T-B1, that can specifically and stably bind thrombin in serum [22], while Tan et al. constructed a circular bivalent aptamer inspired by the circular structure of the enzyme, which greatly improved the stability of the aptamer in nucleases and serum [23].

Although cyclization can stabilize aptamers and enhance their ability to resist nucleases in a simple and rapid manner, it may change the 3D structures of the original linear aptamers and affect their site of action because the current leading method of purifying circular aptamers is PAGE [24]. The recovery rate is low and takes a long time, and the cyclase is relatively expensive; therefore, preparing a large amount of circular aptamers is expensive and time-consuming. Reckless circularization of linear aptamers not only hampers the production of higher-performance aptamers, but may also result in greater economic and time losses. Therefore, there is an urgent need for a method for the rational design of circular aptamers to better assess their impact after cyclization. To the best of our knowledge, our study is the first to propose a more rational approach for designing circular aptamers using molecular computation.

In this study, we evaluate the effect of cyclization of the high-affinity G-quadruplex aptamer 45e1 (5′-CTCGGGGGCGCGGTTGATCGGAGAGGG-3′) targeting STX obtained from Zhou’s SELEX [25], which is cyclized as c45e1. An oxDNA-based circular DNA model is used to construct the 3D structure of the single-stranded circular DNA aptamer c45e1 [26] and 3D-NUS [27] for modelling the G-quadruplex aptamer 45e1. Molecular docking and dynamics are used to evaluate the stability and affinity of the aptamer after cyclization. The combination of molecular computations allows for the reduction of complex upfront activity validation procedures and enables a more standardized and controlled design of circular aptamers, greatly speeding up the development and design of circular aptamers, and finally, promoting their application in therapy.

## 2. Results

### 2.1 Stability analysis of the 3D structures of c45e1 and 45e1

G-quadruplex sequences analyzed using QGRS Mapper [28] for 45e1 are shown in Table S1. G-quadruplex structures modeled with 3D-nus and the 5′ and 3′ ends were extended and patched using Discovery Studio (Ver. 4.5) [29], all visualized with PyMOL (Ver. 2.2.0) [30] shown in S1 a-d, and circular 45e1 modeled with OxDNA shown in S2.

The results of the molecular dynamics simulations (Fig. 1a) showed that the RMSD of 45e1 was consistently higher than that of c45e1 during the 100 ns simulations, indicating a smaller change in the c45e1 conformation. This demonstrates that c45e1 has a more stable structure than 45e1, which is in line with our basic expectations from the circular structure and promises a stable state in a complex environment of body fluids similar to an in vitro environment. The more stable structure of c45e1 may be related to the fact that c45e1 forms more intramolecular hydrogen bonds and dispersed intramolecular hydrogen bonds, resulting in a relatively stable structure. As shown in Fig. 1b, the linear 45e1 hydrogen bonds are only concentrated in the G-core, whereas c45e1 forms multiple knots owing to solvent action and ring tension, making it easier for the individual bases within the ring to form hydrogen bonds in various parts of the ring than in 45e1.

**Fig. 1.**
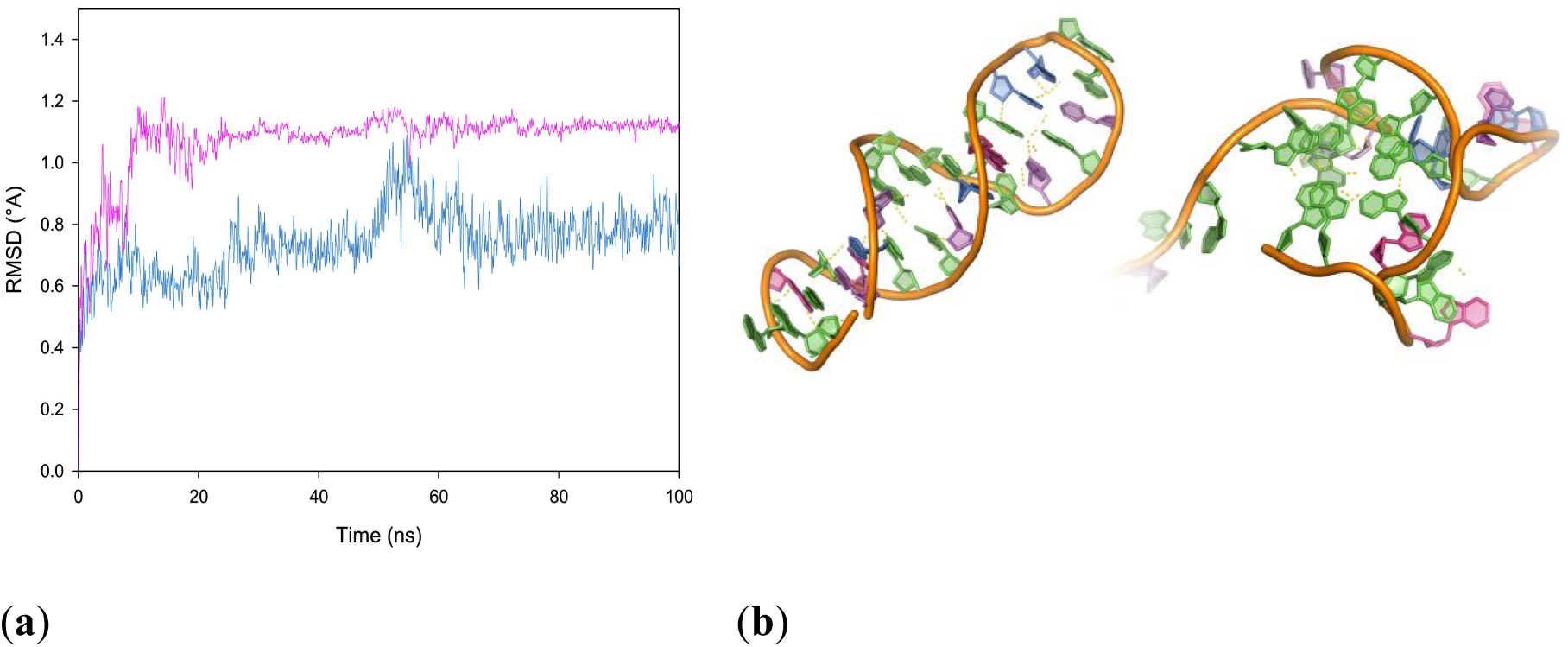
Stability of c45e1 and 45e1 structures. (a) Variation of RMSD for 1000-ns molecular dynamics simulations of 45e1 and c45e1 (purple curve for 45e1, blue curve for c45e1). (b) Distribution of intramolecular hydrogen bonds between c45e1 and 45e1 (c45e1 on the left, 45e1 on the right, yellow dashed lines show the hydrogen bonds formed within the formed molecule, red bases are A, blue bases are T, green bases are G and pink bases are C)

### 2.3. Molecular docking analysis of c45e1 and 45e1 with STX

The molecular docking results of AutoDock Vina 1.2.0 [31] showed that 45e1 binds STX with a binding energy of-7.5 kcal/mol. As shown in Figs. 2a and 2b, there is only one binding site for interaction with STX, and one hydrogen bond is formed. The docking site is G13 on the G-plane, where the epoxy group of the pentasaccharide in G13 interacts with the guanidinium group in STX to form a hydrogen bond. The two G-planes formed by the G-quadruplex (G6, G7, G12, G13, G20, G21, G25, and G26) form a groove-like structure in 45e1, allowing the convex part of the STX structure (the guanidinium group) to be better embedded in the G plane of the quadruplex. This physical binding also stabilizes the complex formed by 45e1 with STX, and the guanidine group is essential for STX toxicity [32], demonstrating that linear 45e1 has the potential to inhibit STX activity.

**Fig. 2.**
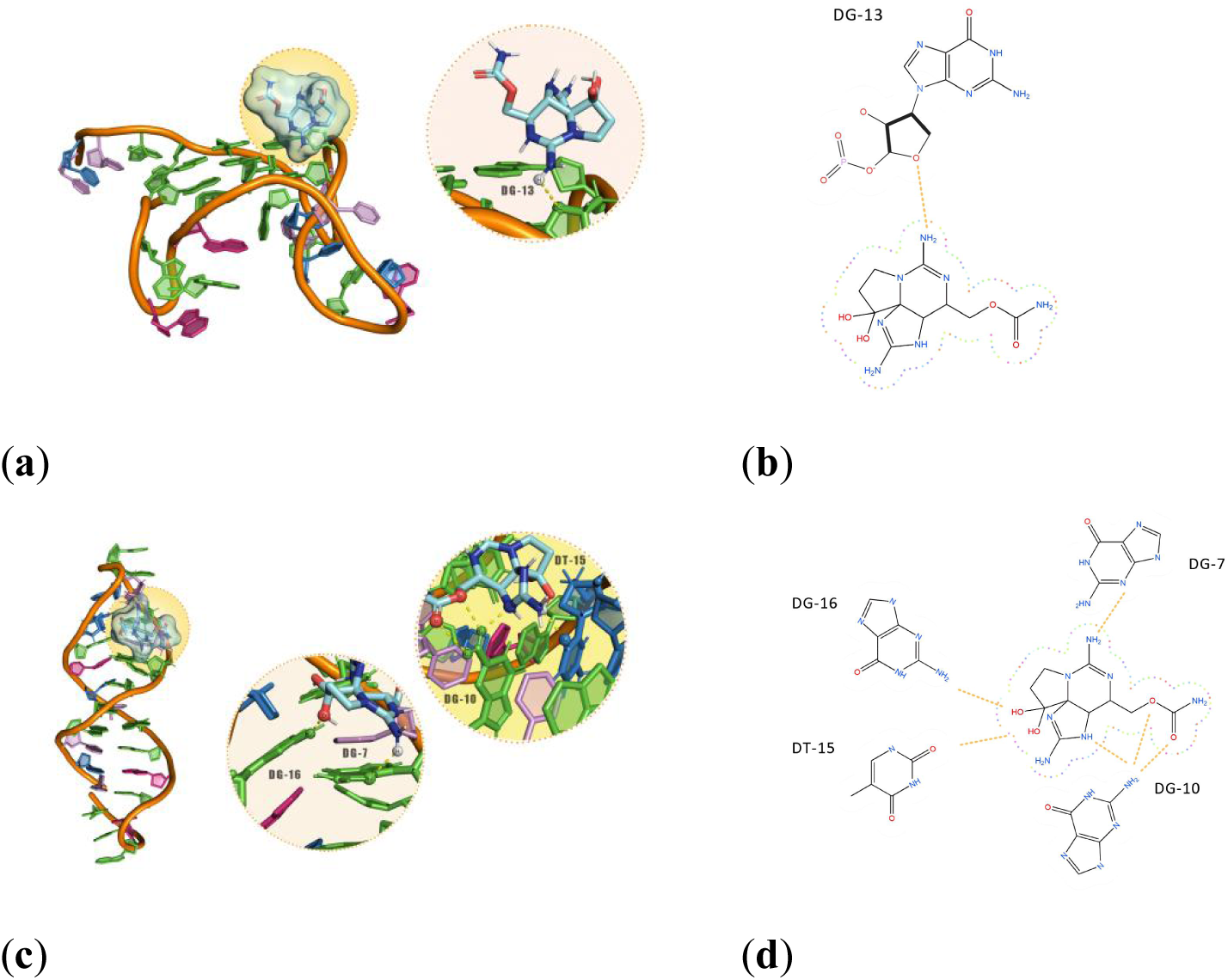
Molecular docking results of c45e1 and 45e1 with STX. (a) 45e1 and STX form a 3D complex structure. (b) 2D interactions between 45e1 and STX. (c) c45e1 and STX form a 3D complex structure. (d) 2D interactions between c45e1 and STX. The yellow dashed lines in (a) and (c), and the light orange dashed lines in (b) and (d) indicate the hydrogen bonds formed by docking.

The binding energy of c45e1 to STX is-7.7 kcal/mol, which means that c45e1 has a higher affinity than 45e1. As shown in Figs. 2c and 2d, c45e1 has more binding sites with STX, forming a total of six hydrogen bonds. G7 forms a hydrogen bond with the guanidine group of STX, G10 forms two hydrogen bonds with the amide group of STX and one hydrogen bond with its pyrrole group, and T15 and G16 form two hydrogen bonds with the two hydroxyl groups. c45e1 binds not only the guanidine group, but also two hydroxyl groups related to toxicity, which may better neutralize STX than 45e1. This makes the structure of the complex more stable than that of 45e1. The cyclization of 45e1 did not affect its binding with STX; rather, it had a higher binding energy and was able to enclose more toxicity-related groups. Thus, compared to 45e1, c45e1 not only has potential for diagnostic applications, but also for the development of applications related to therapy for STX intoxication as it has a stable 3D structure and the ability to resist degradation. However, owing to the high flexibility of the nucleic acid aptamer, molecular docking is more likely to yield a false-positive result [33,34], Further analysis of the molecular dynamics is still required to reveal more realistic and detailed information regarding the interaction of the two aptamers.

### 2.4. Spontaneous docking in molecular dynamics

#### 2.4.1. RMSD trajectory and flexibility analysis during 1000ns spontaneous docking

At 1000 ns (Figs. 3a and 3b), we found that the RMSD of c45e1 was <1 and less than that of 45e1 throughout the spontaneous binding process, indicating that the c45e1 structure is more stable and can form a more stable complex with STX throughout the interaction, in line with our prediction of a more stable circular structure.

**Fig. 3.**
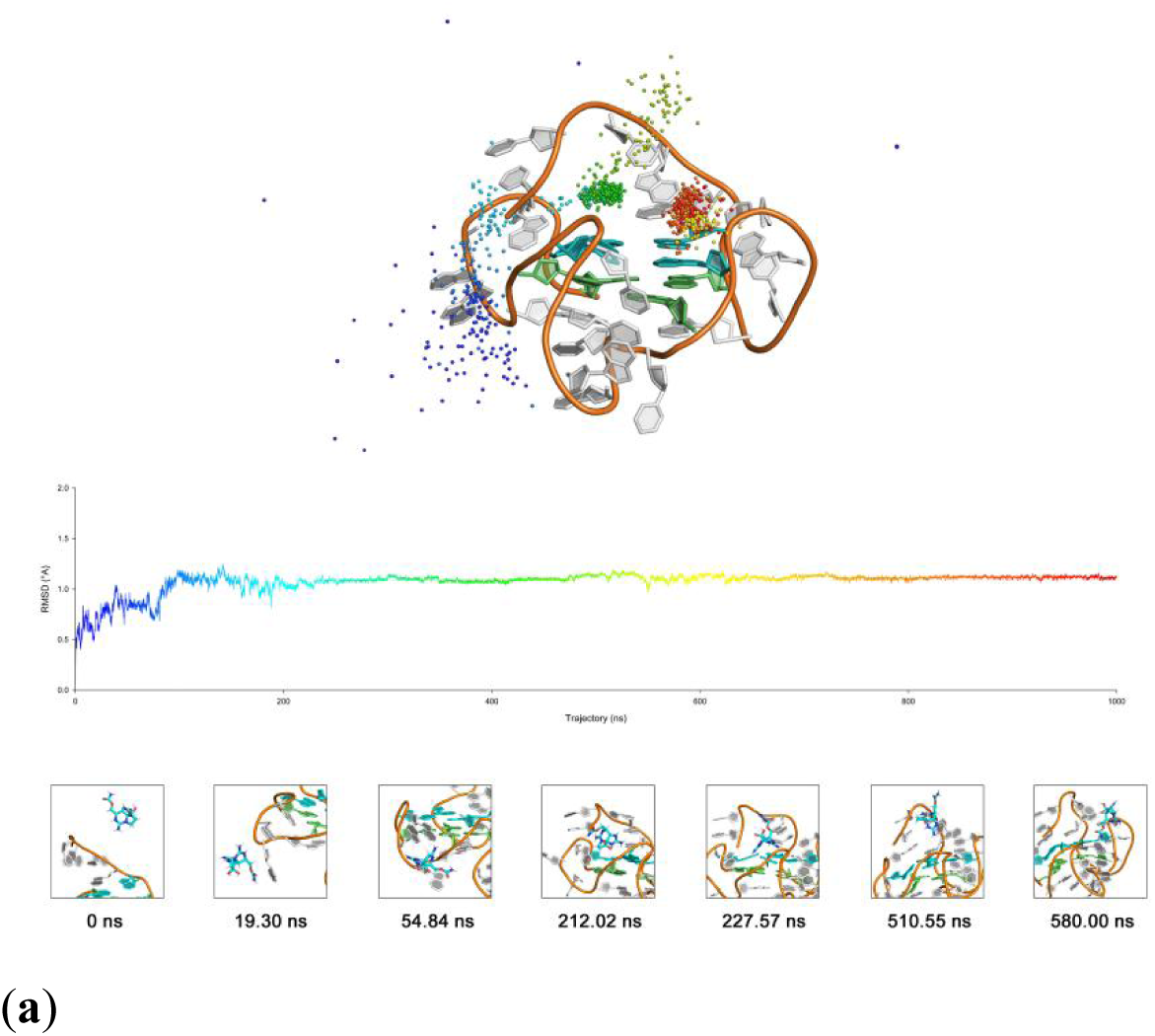
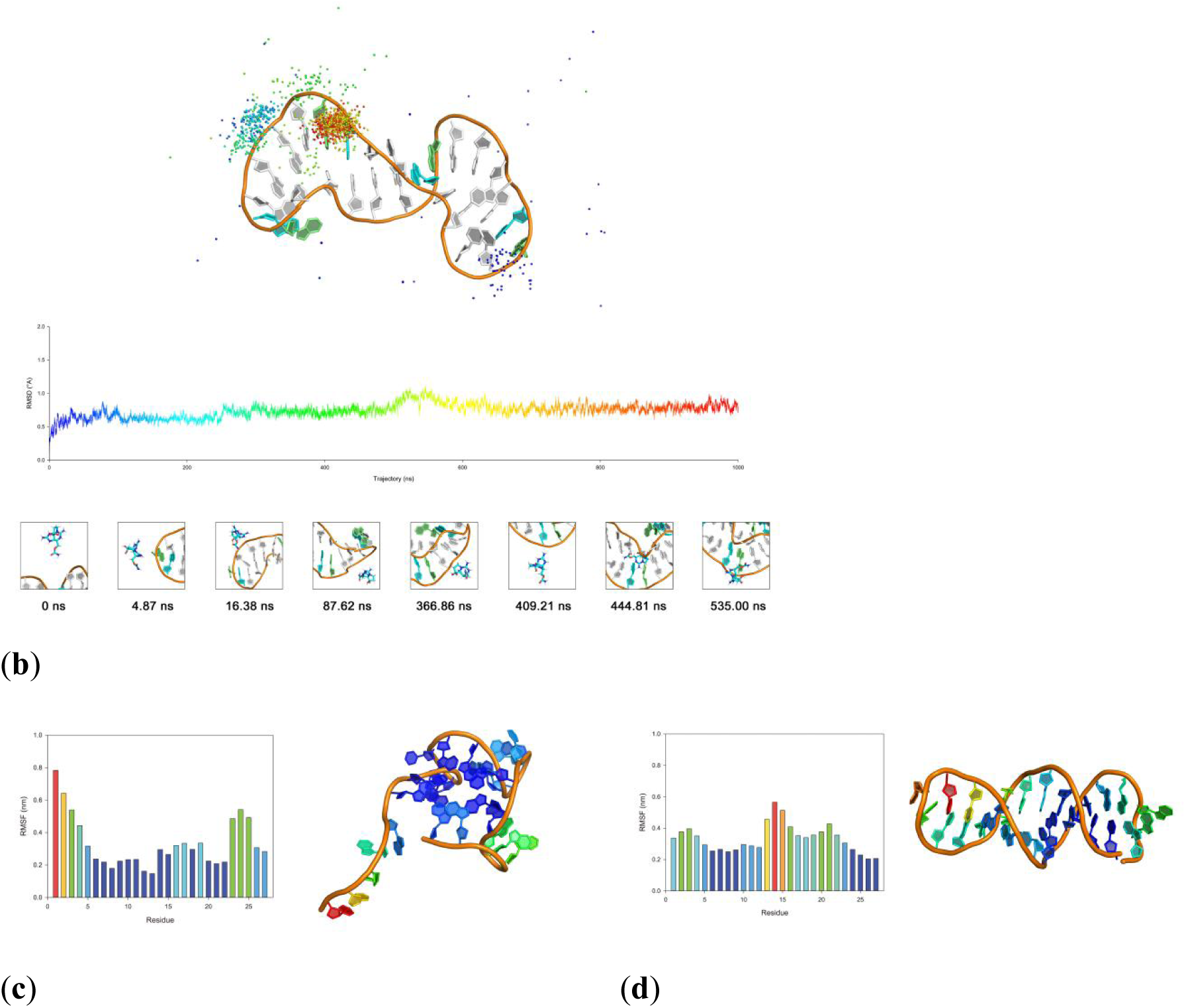
Trajectory and flexibility analysis of 1000-ns spontaneous docking. Molecular motion trajectories of STX during spontaneous docking of (a) 45e1 with STX and (b) c45e1 with STX (different colored spheres represent the change in the center of mass of STX with time, colors are consistent with the curve, and the trajectory of STX is saved once per ns). RMSF of individual bases of (c) 45e1 and (d) c45e1 during spontaneous docking (RMSF bar colors on the left correspond to the residue color in the 3D structure on the right).

For 45e1 (Fig. 3a), STX first moved from a random position 15 nm from the aptamer and contacted 45e1 at 19.30 ns, forming the first hydrogen bond with the phosphate group of A24. It began to approach the G-planes at 54.84 ns and at 212.02 ns it interacted with G20 and G21 in 45e1. Subsequently, STX was embedded in the G-plane at 510.55 ns and interacted with C3 to form a complex similar to that seen above after molecular docking. At 580 ns, STX slipped to the edge of the G-planes and bound stably to G12, finally reaching a stable state.

For c45e1 (Fig. 3b), STX first interacted with G13 of c45e1 at 4.87 ns. It then began interacting with T15, gradually approaching the cavity of c45e1, which is consistent with the docking results. However, it did not remain stable in this cavity and entered a free state at 16.38 ns away from the cavity. At 87.62 ns, it interacted with G23, transitioned to a free state, and gradually became stable. At 366.86 ns it gradually deviated from G23 and began to approach G20 and G21, which are the G-core sequences involved in the formation of the G-planes in 45e1. It first bound to 19C at 409.21 ns, and at 444.81 ns to G4, which is closer to the positions of G20 and G21. Finally, at 535 ns, it stably bound to G20, G21, and A22 to form a stable complex structure. Overall, c45e1 reached a steady state earlier than 45e1, with most of the trajectories clustered around the critical G20 and G21 sites, while some of the trajectories in 45e1 were clustered around the 5′ and 3′ ends owing to interference from them, and on the G-plane where G6, G12, G20, and G26 are located, ending up mostly around G20.

In terms of the overall flexibility of c45e1 and 45e1 (Figs. 3c and 3d), the overall Root Mean Square Fluctuations (RMSF)trend for c45e1 was completely different from that for 45e1. The RMSF of the bases near the 5’ and 3′ ends of 45e1 was significantly higher due to the both ends being in a free state. In contrast, the bases near the 5’ and 3′ ends of c45e1 were more stable, while the less stable sequences concentrated in the middle of 13G, 14T, and 15T, which have significantly higher RMSF. The main reason for this is that cyclization destabilizes the G-quadruplex structure, leading to an increase in the flexibility of the intermediate sequence, and G-plane bases in the vicinity are likely to be able to escape from the Hoogsteen hydrogen bonding network. This makes the G20 site of c45e1 different from that of 45e1, and the greater flexibility allows more bases around G20 to be involved in the interaction with STX, generating more potential binding sites. The overall trajectory analysis clearly suggests that perhaps c45e1 has more potential to treat the toxin, being able to bind it earlier and for longer. Not only are its own 5’ and 3′ ends not in a free state, but also, being less flexible, they do not bind STX and prevent it from binding to the critical binding site, revealing that G20 has similar roles in both c45e1 and 45e1 as the base that primarily binds STX. This also suggests that the two aptamers may have similar modes of interaction, and that the cyclization of 45e1 may not affect binding. Furthermore, the disruption of the G-quadruplex structure increased the flexibility of the G-plane bases, allowing c45e1 to have more bases involved in the interaction than 45e1. This suggests that c45e1 may bind to STX more firmly.

#### 2.4.2. Analysis of the binding situation during 1000-ns spontaneous docking

To quantify the extent of binding between the two aptamers, we plotted the binding-energy landscape. The reaction of the two aptamers with STX throughout the process is indicated by the size and depth of the high-energy areas (Figs. 4a–4d). The binding energy landscape of 45e1 showed two energy valleys, similar to that of c45e1; however, one of the energy valleys had a significantly lower energy depth and area than c45e1, suggesting that c45e1 may have a better ability to bind STX.

**Fig. 4.**
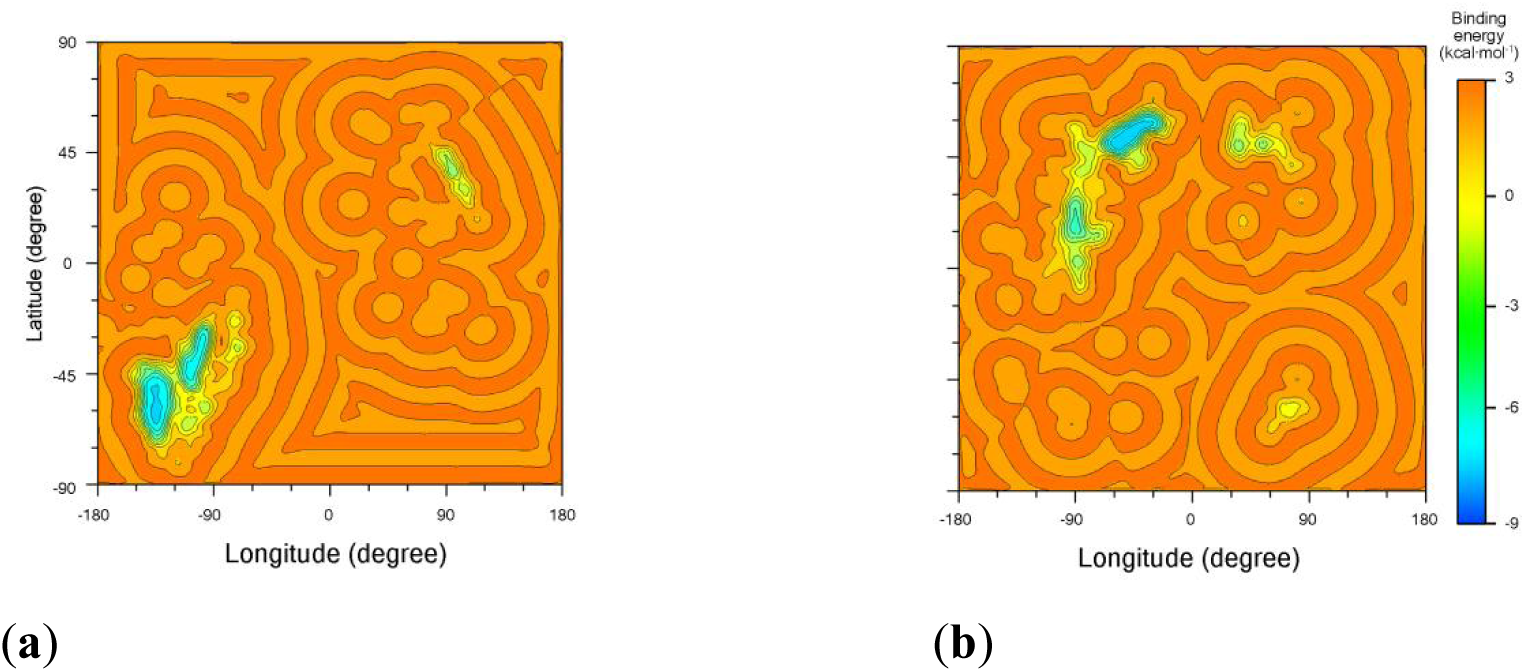
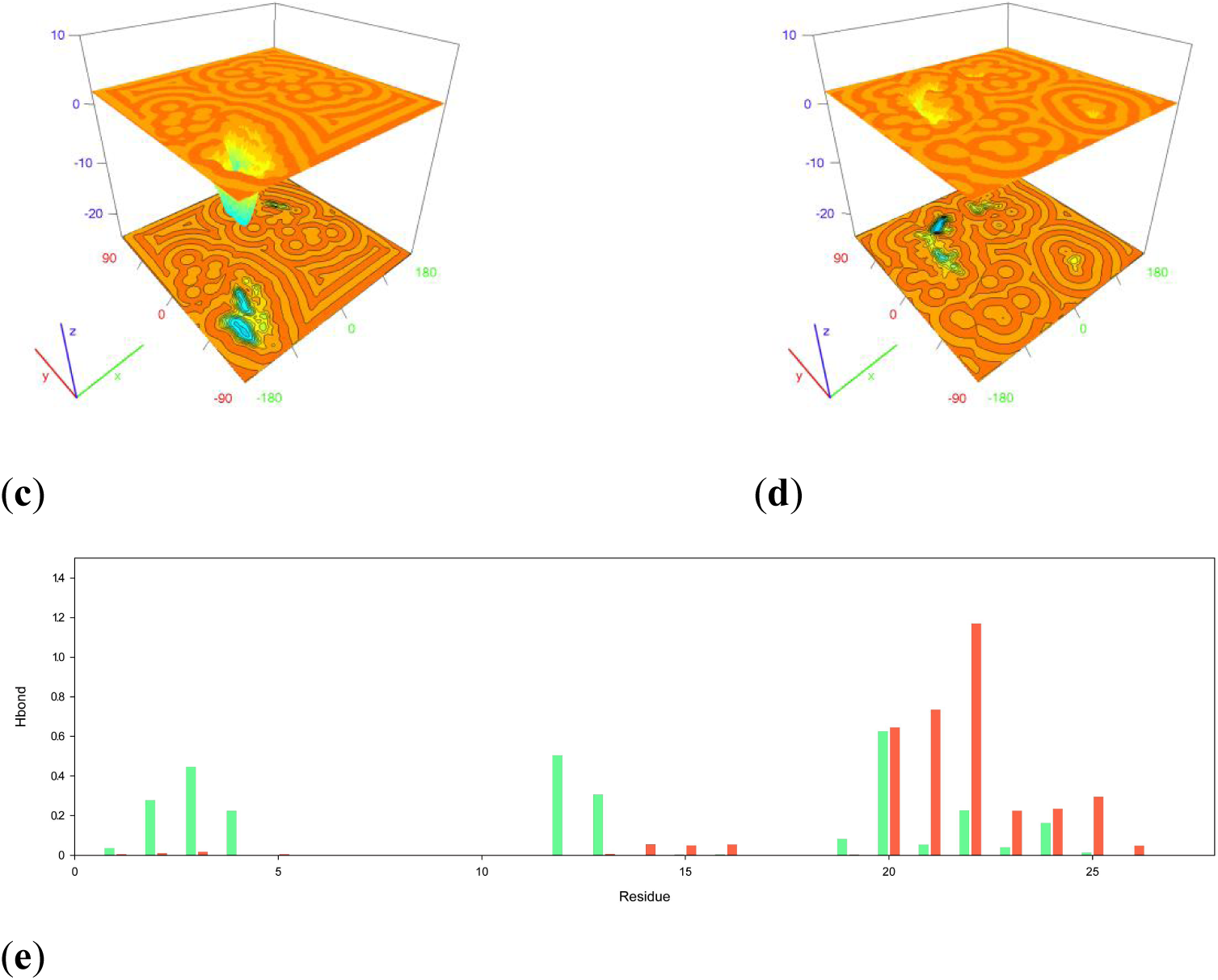
Binding of STX to 45e1 and c45e1 in 1000 ns. 2D binding energy landscape of the interaction between (a) 45e1 and STX, and (b) c45e1 and STX during spontaneous docking. 3D binding energy landscape of the interaction between (c) 45e1 and STX, and (d) c45e1 and STX during spontaneous docking. (e) The number of hydrogen bonds of each residue per ps in c45e1 and 45e1 during spontaneous docking (red column for c45e1, blue for 45e1).

The number of hydrogen bonds formed by each base also shows that there are some differences in the number and distribution of bonds between the two aptamers (Fig. 4e). The hydrogen bonds formed by c45e1 are mainly concentrated in G20 and its surrounding bases G21, A22, G23, A24, and G25, whereas those formed by 45e1 are distributed among the sequences associated with the formation of the G-quadruplex (G12, G13, G20, G21, A22, G23, and A24) and some of the bases at the 5′ end (T2, C3, and G4), where G20 is found in both 45e1 and c45e1, forming more hydrogen bonds and playing a more critical role. The main reason for this difference in hydrogen bond distribution is that, unlike 45e1, the G-plane in c45e1 is disrupted, causing bases such as G12 and G13 to move away from the G-plane sequence associated with G20 and resulting in the hydrogen bonds being concentrated mainly on the bases around G20. Second, the 5′ and 3′ ends of c45e1 are not in a free state, which allows the key site to be more easily exposed and reduces obstruction from the 5′ and 3′ ends, allowing STX to bind to the key site earlier, so that fewer hydrogen bonds are formed at other sites. In contrast, due to the impact of free fragments at the 5′ and 3′ ends and the G-quarters in 45e1, the hydrogen bonds are distributed over the critical G20 and its surrounding G-quadruplex sequences as well as the 5′-end free fragment, which also prevents the toxin from binding to the G20 site.

Molecular dynamics revealed the specific effects of cyclization on 45e1 in terms of molecular properties before and after cyclization, as well as the binding ability. It confirmed that c45e1 may have higher performance than 45e1, bind toxins earlier, and form a more stable complex, not only sharing a key site of action with 45e1, but also exhibiting higher affinity potential. These results suggest that although the original G-quadruplex structure of c45e1 is disrupted by cyclization, the key site of action is retained, and c45e1 retains its original high affinity and is valuable for mass cyclization.

### 2.4. Validation of the affinity of c45e1 for STX by Bio-layer interferometry(BLI)

The cyclization of 45e1 and the associated large-scale purification system are illustrated in S2 and S3. By immobilizing the biotin-labelled c45e1 obtained earlier and interacting it with STX, the K_on_ value of c45e1 binding to STX was found to be 4.31 x 10^4^ 1/Ms, the K_dis_ value was 7.31 x 10^-4^ 1/s, and the Kd value was 1.70 x 10^-8^ M. The affinity constant of c45e1 is less than that of 45e1, as measured by Zhou using BLI (1.90 x 10^-8^ M)[25]. This indicates that not only can it maintain a similar affinity as that of 45e1, the affinity is slightly improved after cyclization, demonstrating that the 3D structure of c45e1 predicted by OxDNA does have certain accuracy. The molecular computation pipeline also provides a more accurate analysis and prediction of the effects of 45e1 cyclization, which is fully consistent with the trends obtained from the experimental analysis. Table 2 lists the specific parameters obtained from the BLI validation of c45e1.

**Table 1.** BLI results of c45e1.

| Type of sensor | Sample | K<sub>d</sub> (M) | K<sub>d</sub> Error | K<sub>on</sub> (1/s) | K<sub>dis</sub> (1/s) | K<sub>dis</sub> (1/s) |
| --- | --- | --- | --- | --- | --- | --- |
| Streptomycin | c45e1 | 1.70×10<sup>-8</sup> | 1.02×10<sup>-9</sup> | 5.04×10<sup>-3</sup> | 7.31×10<sup>-4</sup> | 3.29×10<sup>-5</sup> |

K<sub>d</sub>: affinity constant, K<sub>dis</sub>: dissociation rate constant, K<sub>on</sub>: binding rate constant

**Table 2.**
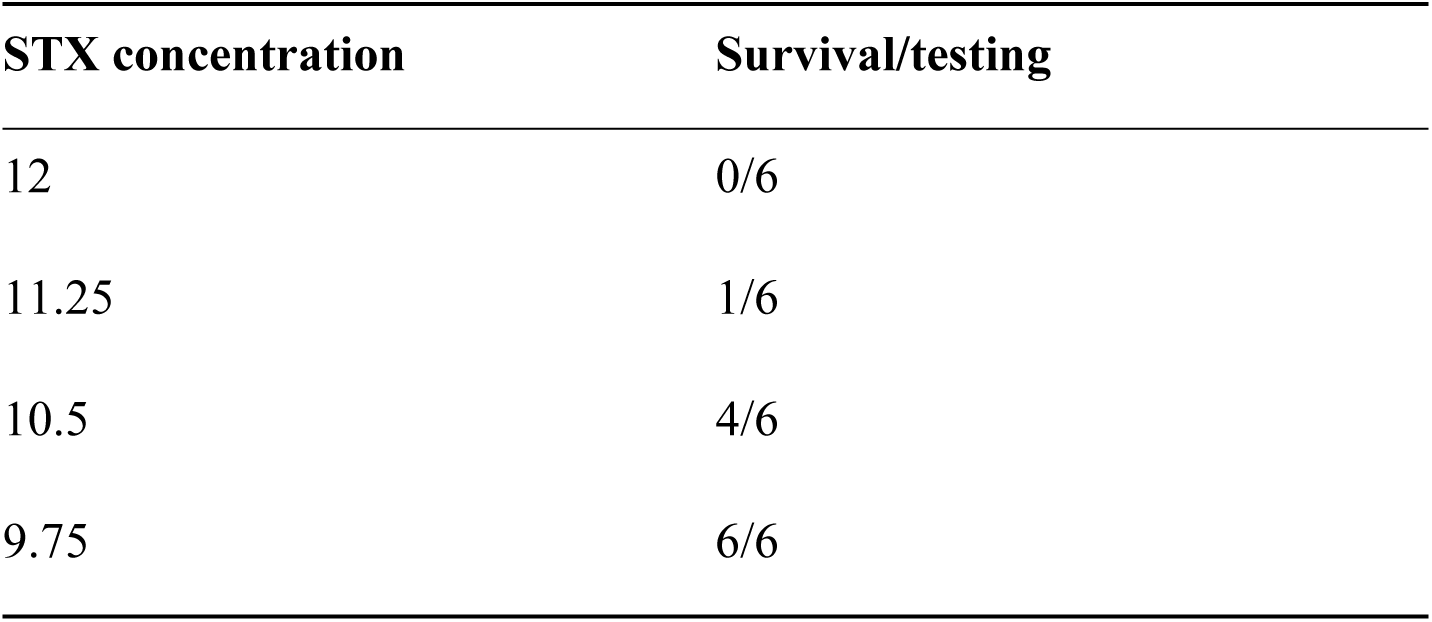
Relative toxicity of STX in mice at different doses.

### 2.5 Circular dichroism(CD) spectrum to verify aptamer structures

Initially, as shown in Figure 5A, the negative ellipticity at 240 nm and the positive ellipticity at ∼280 nm unambiguously indicate that 45e-1 adopts a parallel G-quadruplex topology. Comparison of the CD spectra of 45e-1 before and after incubation with STX (Figure 8C) reveals negligible spectral changes, demonstrating that STX binding does not perturb the G4 topology, which remains highly stable under these conditions.

**Fig. 5.**
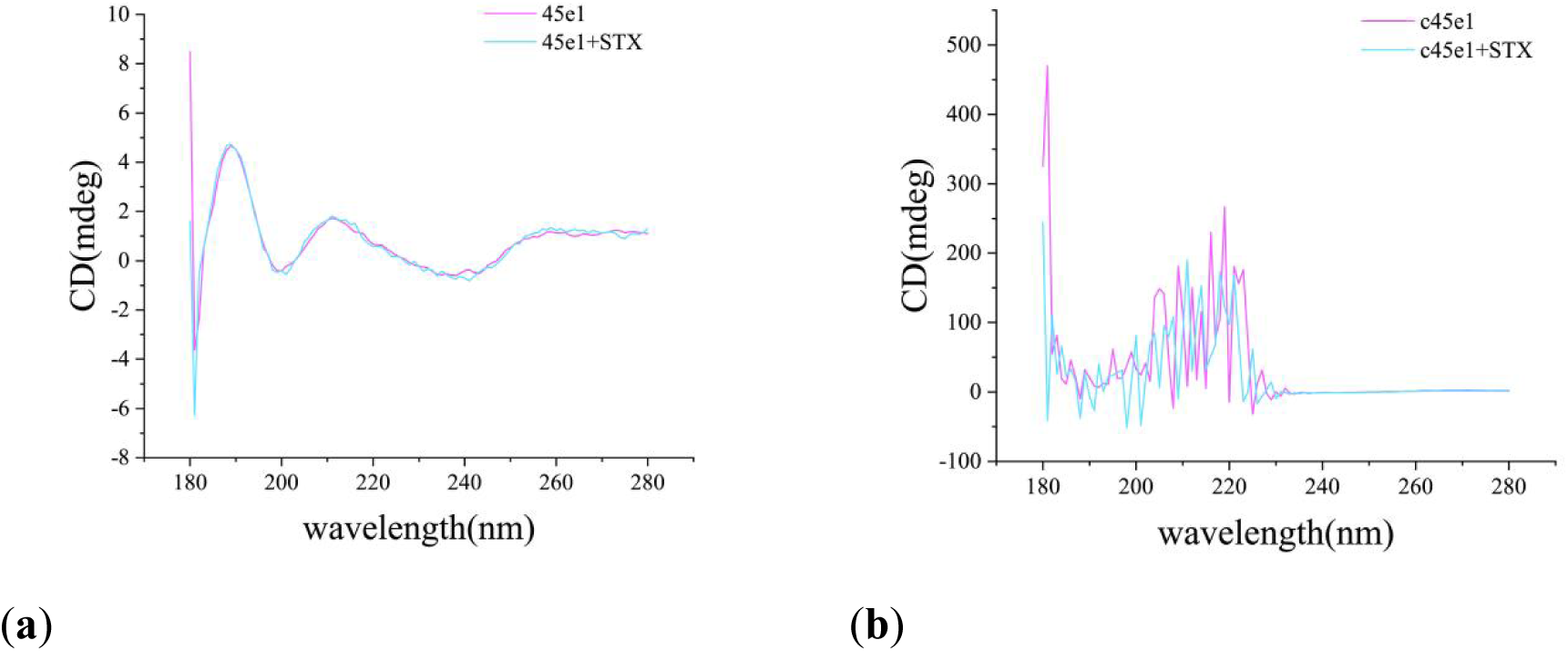
CD spectra of aptamer 45e1 and c45e1 before and after binding to STX in 6uM. (a) Pure 45e1 and 45e1-STX. (b) Pure c45e1 and c45e1-STX.

**Fig. 5.**
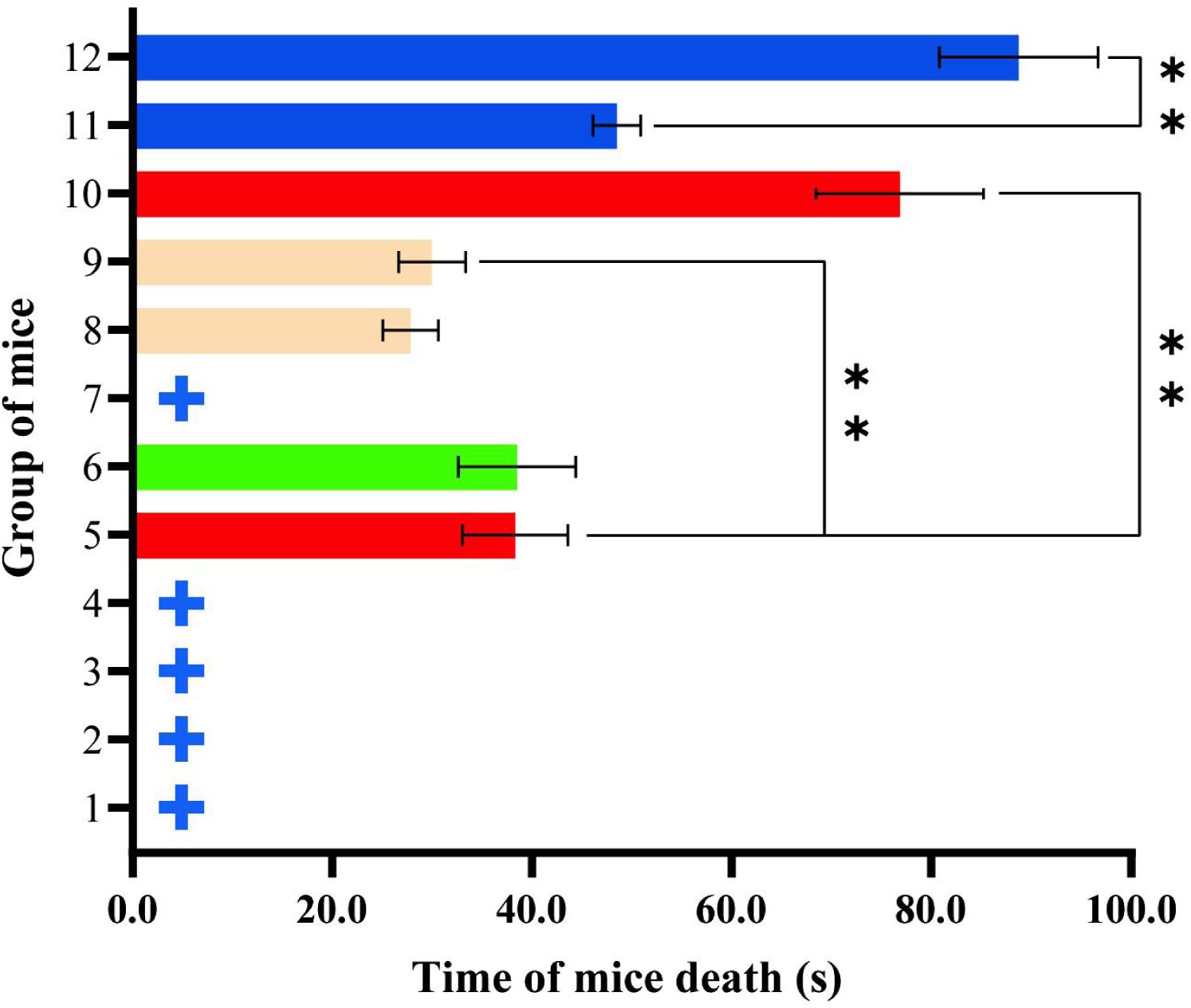
Grouping results of caudal venous injection experiment in mice. 1: Linear aptamer; 2: cyclization reaction mixture of aptamer; 3: cyclization reaction mixture after passing through 10-kD ultrafiltration membrane; 4: Circular aptamer after purification; 5: STX (11.25 μg/kg); 6: STX (11.25 μg/kg) + linear aptamer 45e1; 7: STX complex (11.25 μg/kg) and linear aptamer passed through 3-kD ultrafiltration membrane; 8: STX + cyclization reaction mixture of aptamer; 9: STX (11.25 μg/kg) + cyclization reaction mixture of aptamer through 10-kD ultrafiltration membrane; 10: STX (11.25 μg/kg) + circular aptamer after purification; 11: STX (11 μg/kg); 12: STX (11 μg/kg) + circular aptamer after purification; +: surviving mice (six). The error bars represent standard error (n = 3), **p < 0.01.

Subsequently, CD spectra of the cyclic aptamer c45e-1 were recorded at 6 μM, 2 μM, 0.67 μM, 0.22 μM, and 0.07 μM (Figure S4). In a properly executed dilution series, one expects a systematic decrease in peak intensity accompanied by smooth absorption profiles. Therefore, the sharp features observed below 230 nm in Figure 5B are attributed to instrumental noise, and no dichroic signal is detected above 230 nm. This absence of CD activity confirms that c45e-1 lacks chiroptical signatures of a G-quadruplex, implying that cyclization disrupts the native quadruplex topology. These results are fully consistent with prior computational predictions, which forecast that cyclization of 45e-1 would abolish its original G4 conformation and yield a symmetric, non-chiral structure.

### 2.6. Experimental assessment of the safety and efficacy of circular aptamers in mice

Four STX concentration gradients were established for this experiment. The results of this experiment are consistent with those of previous studies, but differ significantly from the data reported in the literature (LD_50_=3.4 μg/kg) [35]. The results of STX toxicity in mice are shown in Table 3; the final concentration of STX was selected to be 11.25 μg/kg for groups 1–10, while groups 11 and 12 were treated with a concentration of 11 μg/kg.

The interaction between the aptamer and STX was evaluated in a multi-group controlled experiment, and the results of aptamer safety and efficacy validation are shown in Fig. 5. Groups 1, 2, 3, and 4 showed no significant toxicity in mice for any of the components involved in the experiments. The results of Group 6 showed that there was no significant difference in the time to death between mice injected with the linear aptamer 45e1 mixed with STX and mice injected with STX alone (Group 5), although 45e1 could bind with high affinity to STX in vitro and had no protective effect on mice in vivo. This suggests that 45e1 is rapidly degraded in vivo and thus does not exert a protective effect. As shown in the results of Group 7 in Fig. 5, the mixture of 45e1 and STX was filtered through a 3kD ultrafiltration membrane and then injected intravenously into mice, all of which survived. Due to the inability of the toxin-aptamer complex to pass through the ultrafiltration membrane, the actual concentration of STX injected into the mice was much lower than the lethal amount in the experimental data, which proves that 45e1 binds to STX but does not prove that it inhibits the biological activity of STX.

As shown in Group 10 of Fig. 5, the mice were injected with the circular and purified aptamer c45e1 mixed with STX. Although all mice eventually died, the mean time to death was 2-fold that of mice injected with STX alone (group 5), with a significant difference in the time to death between the two groups on one-way analysis of variance (p < 0.01). This suggests that c45e1 exerts a protective effect in vivo. After reducing the toxin concentration, as shown in experimental groups 11 and 12, the survival rate of the mice did not improve, but the time to death was significantly longer (p < 0.01). This may be due to the small molecular weight of the circular aptamer (8.8 kD), which is easily filtered in vivo. With reference to previously marketed aptamer drugs, modifications such as PEGylation of the aptamer may prolong the half-life of the aptamer or increase its stability in vivo [36], which can be further optimized in subsequent experiments. Another reason may be that the binding of the aptamer to STX remains in dynamic equilibrium after entering mice; therefore, STX toxicity cannot be completely inhibited.

## 3. Conclusion and Discussion

In this study, we established a pipeline to predict the 3D structure of circular aptamers and built a molecular computation pipeline to better evaluate the rationality of circularization. Simultaneously, we validated the performance of our model and predicted its structure using BLI and mouse experiments. Our results from molecular computation revealed that cyclization disrupts the stable G4 structure allowing the unconstrained G4 sequences to become more flexible, and the 5’ and 3’ ends to not be in a free state, which prevents the 5’ and 3’ free fragments from impeding the binding with STX. Meanwhile the flexible G4 sequence provides more bases and extra space to better capture STX. This eventually leads to c45e1 not only having higher stability during the simulation process, but also capturing STX earlier and having a higher affinity than 45e1. The final BLI and mouse experiments confirmed that cyclization did not affect the binding ability of c45e1; similar to the trend predicted by simulations. c45e1 showed a protective effect, increased affinity, and significantly prolonged the survival time of mice, suggesting that the circular aptamer designed and validated by this method is not only efficient but also reliable.

Because of the difficulty in obtaining the 3D structures of aptamers experimentally, the mechanism of action of most aptamers is unknown [37], making computational methods an indispensable approach to obtaining their 3D structures, analyzing their mechanisms, and designing them. Therefore, researchers have used a variety of computational methods to design linear aptamers for a wide range of targets [38,39,40] and there are no specific procedures for analyzing and designing circular aptamers. Owing to the excellent activity and anti-degradation ability of some linear aptamers after cyclization, an increasing number of circular aptamers have been emphasized by researchers for their potential as therapeutic drugs [24,41,42]. Therefore, a method to efficiently design and evaluate circular aptamers can better promote the development of such aptamer-based drugs. As mentioned earlier, the new method for the design and evaluation of circular aptamers proposed here can efficiently confirm whether some linear aptamers are unaffected by cyclization. It can be used to rationally design circular aptamers and obtain atomic-level dynamic information for analyzing their mechanisms, which adds more channels for improvement.

In summary, this study successfully evaluated the effect of 45e1 after cyclization and designed a high-affinity cyclic aptamer that may have potential applications in the development of STX therapeutic drugs. In addition, our rational design of the circular aptamer suggests that the new approach is promising for the design and exploration of mechanisms to optimize the circular structure for subsequent drug ability assessments.

## 4. Materials and Methods

### 4.1. Building the 3D Structure of 45e1

The sequence of 45e1 is (5’-CTCGGGGGCGCGGTTGATCGGAGAGGG-3’), which has a guanine-rich portion and at least four pairs of discontinuous guanines, indicating that there is a high probability of forming G-quadruplex in this sequence. According to the modelling method by Song et al. [43], the 45e1 sequence was entered into the QGRS Mapper to analyze the input sequence for the presence of fragments that could form a G-quadruplex. The high-scoring fragments from QGRS Mapper were then input into 3D-Nus to construct all nine possible types of conformations of Gcore sequences in 45e1. Finally, the sequences around the 5’ and 3’ ends was extended using the build/edit nucleic acid function in Discovery studio (Ver. 4.5) by selecting create/grow at 5′ and 3′ in DNA. The 5′ and 3′ extended nucleic acid fragments were then ligated using the ligate function. Missing atoms or broken fragments were added to the full-length sequence, and the broken phosphodiester bonds of the gaps were ligated. Finally, nine possible G-quadruplex structures of 45e1 were constructed, and the stability of the nine structures was investigated through temperature-dependent simulations of molecular dynamics. The structure with the lowest RMSD fluctuation was selected as the most likely natural 3D structure of 45e1 and used for further verification.

### 4.2. Building the 3D Structure of c45e1

First, a new input file was created for adjusting basic parameters, and temperature was set at 20 °C with a salt concentration of 0.5 M and steps set to 10^9^. The trajectory file named traj.dat was exported. Next, the topology of the aptamer nucleotide sequence targeting the biotoxin was imported into the experimental environment for a virtual Monte Carlo dynamics simulation to achieve rotation, translation, and movement of the nucleotide [44], creating trajectory data for steps 1–10^9^. The final fitted trajectory data at different threads containing energy information were exported and then sorted according to the value of the total energy (E_total_), identifying the trajectory with the lowest E_total_ for all 10^9^ steps. However, since the trajectory calculated and folded by OxDNA is a coarse-grained model, it was difficult to use it for all-atom high-precision molecular docking and dynamic analysis. We used tacoxDNA [45] software to import the energy-minimum trajectory data and topology, converted the coarse-grained model into an atomic-level model, and exported the pdb file. Note that at this point the 5′ and 3′ ends of the circular structure were still free and not linked by a phosphate diphosphate bond; therefore, the terminal hydrogen atoms of 5′ and 3′ had to be removed from the Discovery Studios and the Sketch Molecules-> Sketch function was used to refine the phosphate group at 5′ to make it bond with the 3′ terminal oxygen atom. The Sketch Molecules-> Clean Geometry function was used to optimize incorrect bond lengths and adjust the global structure.

Meanwhile, the all-atom model derived from tacoxDNA transformation also had chain breaks; therefore, we still needed to use the method described in Section 4.1 to connect the broken parts and replace the missing atoms.

### 4.3. Constant temperature molecular dynamics simulation analysis

Thermostatic molecular dynamics simulations of the full-length aptamer monomer model were performed using GROMACS (Ver. 5.1.4) [46]. Parameters such as the bond angles of nucleic acid aptamers were set using the Amber19bsc1 force field [47]. It is worth noting that conventional nucleic acid structures such as 45e1 top files can be converted by the command pdb2gmx that comes with Gromacs. However, since the circular structure does not have a conventional closed 5′-3′ end in Gromacs, the deletion of the terminal hydrogen atom must be written into the tab file to create a normal gro file, and the bonding information for the atoms associated with the terminal phosphate is written into the generated top file for the next molecular dynamics simulations.

The topology and interaction force parameters of the solvent water molecules were set using the TIP3P water model [48]. The initial position of the aptamer models was then set at the center of the water box, and the distance of the surface of each model from the box boundary was greater than 15 Å. In order to maintain a neutral environment in the molecular dynamics system, a certain number of Mg^2+^ and Cl^-^ ions were placed with a random replacement of the water molecules and the concentration of ions in the solution was made up to 150 mM. The simulations were carried out using periodic boundary conditions, all covalent bonds in the system were constrained by the LINCS algorithm, and the simulation time step was 2.0 fs [49]. The pressure of the NPT system was maintained at 1 atm using a Berendsen thermostat, and the temperature was controlled at 300 K using a velocity-rescaling algorithm [50, 51]. A ramp-up simulation was performed using the simulated annealing method in GROMACS. In the constant temperature phase, the aptamers were simulated at 300 K for 100 ns. The RMSD values for the aptamer monomer were calculated using the command gmx rms, and intramolecular hydrogen bonds were visualized using PyMOL (Ver 2.5).

### 4.4. Molecular docking

First, the STX molecule was downloaded in SDF format from the PubChem (https://pubchem.ncbi.nlm.nih.gov/) website, the correct bond lengths and bond angles in the 3D structure of STX were obtained via clean function in PyMOL, and the file format of STX was converted to PDB format. STX was used as the receptor for molecular docking, and the constructed 45e1 and c45e1 were used as receptors. Subsequently, the lattice-point energy calculation files were created using AutoDock (Ver. 4.2) software. Since we did not determine the exact docking range of 45e1 and c45e1 in the aptamer, we manually adjusted the size of the docking box so that it could encompass the entire aptamer and allow all bases to participate in docking, setting both the lattice spacing and lattice edge length to 0.4 Å. Finally, we saved the software output lattice calculation parameter file in gpf format and ran the AutoDock AutoGrid (Ver. 4.2) plug-in. The docked receptors and ligands were then converted into pdbqt files using AutodockTools 1.5.1. Molecular docking was performed using AutoDock Vina (Ver. 1.2.0) software, where the number of calculations for the genetic algorithm was set to 10, number of populations to 150, maximum number of iterations to 2500000, search algorithm to Lamarckian genetic algorithm, and all other parameters to the default values. The docking parameters were saved as dpf files, and the analysis output as a dlg file. The binding energies of aptamers 45e1 and c45e1 with STX were calculated using the AutoDock Vina software and visualized using PyMOL.

### 4.5. Molecular dynamics simulation of spontaneous binding

This method mainly refers to the spontaneous binding simulation and energy-binding landscape plot by Liu et al [52]. First, the top file for STX was generated using ACPYPE [53] with GAFF and AM1BCC charge paramters[54,55]. The remaining parameters of the spontaneous binding simulations were similar to those in the aforementioned thermostatic simulations. Subsequently, Mg^2+^ was added to the GQ site to maintain the stability of the GQ conformation [43] using a PyMOL script (center of mass). The RMSD values for the spontaneous binding process were obtained using the command gmx rms. Because different software programs have different criteria for hydrogen bonding, only the gmx hbond in GROMACS 5.1.4 was used to calculate the number of hydrogen bonds formed in STX with each base of the two aptamers.

### 4.6. Preparation of c45e1

#### 4.6.1. Small-scale cyclization of linear 45e1

The cyclization reaction system was prepared according to the ratios listed in Table 4. The system was first incubated at the optimum ligase action temperature of 60 °C for 1 h and then the cyclization reaction was terminated by increasing the temperature to 80 °C to inactivate the enzyme. However, owing to the uncertainty of the position of c45e1 on PAGE, a simple pre-experiment was required to determine the relative positions of 45e1 and c45e1 to facilitate the subsequent bulk preparation of c45e1.

**Table 4.**
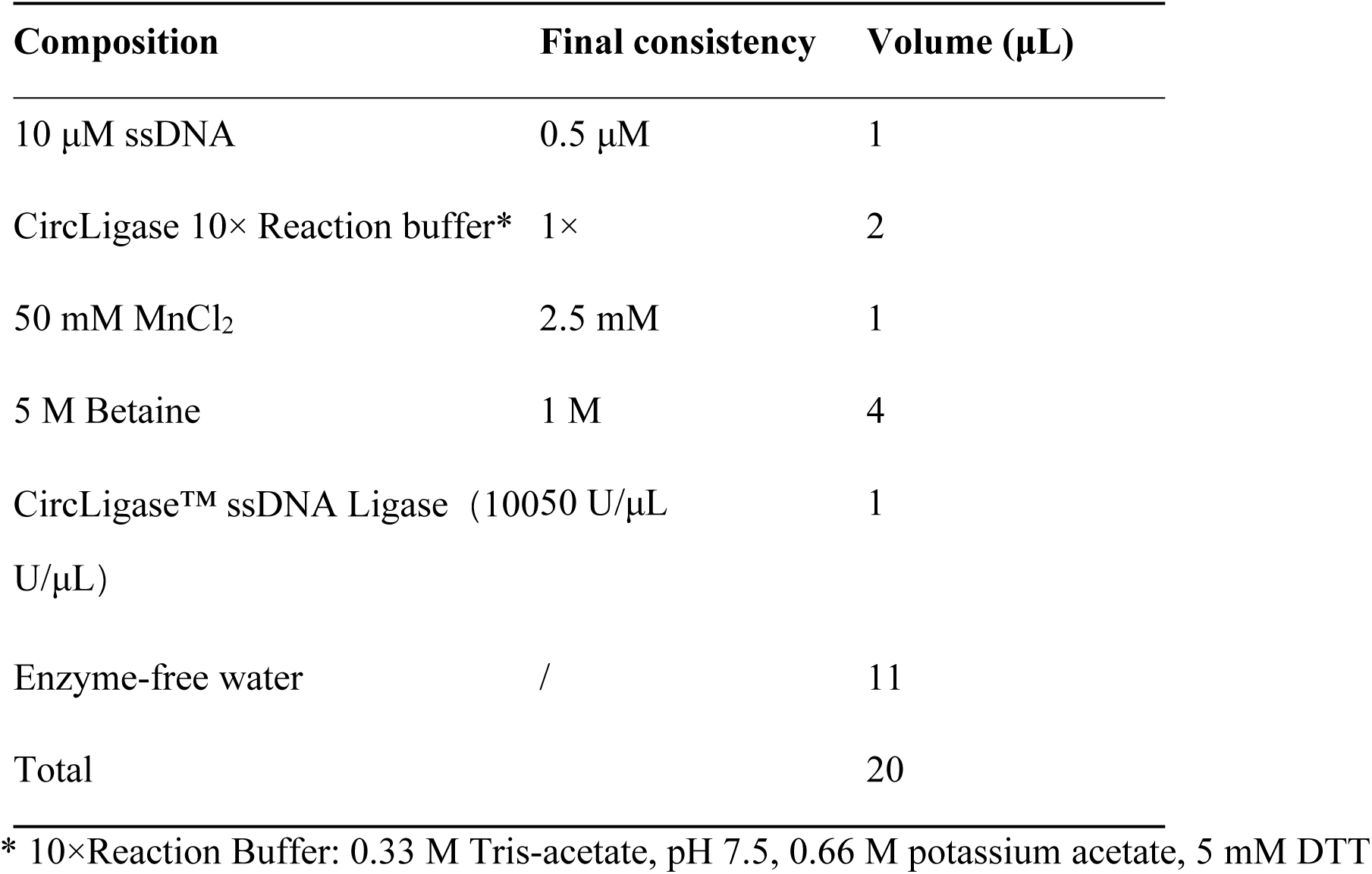
Composition of the cyclization reaction system.

#### 4.6.2. Bulk preparation of c45e1

For the bulk preparation of circular ssDNA for further experiments, a 1.5 mm 15% non-denaturing PAGE gel was used for electrophoresis. The reagent recipe is shown in Table 5 and the constant voltage was set at 300 V for 20–35 min. The gel was then cut for recovery, purified using a Qiagen gel extraction kit, and finally added to the screening buffer for storage.

**Table 5.** Composition of 15% non-denaturing PAGE.

| Composition | Volume |
| --- | --- |
| ddH<sub>2</sub>O | 5.27 mL |
| 10× TBE buffer | 1.20 mL |
| 40% acrylamide | 3.94 mL |
| 10% AP | 72.00 μL |
| TEMED | 18.00 μL |

TEMED: tetramethylethylenediamine, TBE: Tris/Borate/EDTA

### 4.7. BLI validation of circular aptamer affinity with STX

The 5′-end phosphorylated ssDNA 45e1 with a biotin tag (5′P&bio-45e1) was cyclized, and in next stage, the circular ssDNA was recovered and purified via non-denaturing PAGE. The final step of purification was carried out by re-solubilizing the DNA with screening buffer and preserving the reserve. Since the biotin tag is carried on the c45e1 sequence, it can bind firmly to streptavidin, thus coupling c45e1 to the sensor probe and interacting with free STX. The binding rate constant K_on_ and dissociation rate constant K_dis_ were measured, and the affinity constant Kd was calculated.

### 4.8 Circular Dichroism (CD) Spectroscopy Experiments

#### 4.8.1 Preparation of Nucleic Acid Samples by Lyophilization

Owing to the relatively high concentration required for CD measurements in this study, the cyclic aptamer c45e-1 was subjected to pre-concentration. The sample was first held at –80 °C for 24 h, then lyophilized to dryness. The resulting powder was reconstituted in 400 µL of RNase-free H₂ O, and the nucleic acid concentration was determined using a fluorescence-based nucleic acid quantitation instrument.

#### 4.8.2 CD Measurement of Aptamer Conformation

A. The linear aptamer 45e-1 and its cyclic counterpart c45e-1 were each dissolved in RNase-free H₂ O to a final concentration of 12 µM. Solutions were heated at 95 °C for 10 min in a metal bath and then immediately chilled on ice for 5 min. Saxitoxin (STX) was separately dissolved in RNase-free H₂ O and diluted to 12 µM.

B. Six 0.2 mL PCR tubes were labeled 1 through 6 and prepared as follows (all volumes in µL):

Tube 1: 170 µL RNase-free H₂ O (blank control)

Tube 2: 85 µL 12 µM RNase-free H₂ O + 85 µL 12 µM STX (to yield 6 µM STX)

Tube 3: 85 µL 12 µM RNase-free H₂ O + 85 µL 12 µM 45e-1 (to yield 6 µM linear aptamer)

Tube 4: 85 µL 12 µM STX + 85 µL 12 µM 45e-1 (to yield 6 µM each of STX and linear aptamer)

Tube 5: 85 µL 12 µM RNase-free H₂ O + 85 µL 12 µM c45e-1 (to yield 6 µM cyclic aptamer)

Tube 6: 85 µL 12 µM STX + 85 µL 12 µM c45e-1 (to yield 6 µM each of STX and cyclic aptamer)

C. Prior to measurement, each sample was transferred sequentially into a quartz cuvette, rinsing the cuvette with RNase-free H₂ O between samples. Measurements were conducted in an oxygen-depleted sample chamber. If sample signals exceeded the instrument’s linear range, appropriate dilution with RNase-free H₂ O was performed to maintain equal concentrations.

D. Spectral acquisition parameters were set as follows:Wavelength range: 180–280 nm, Bandwidth: 1.0 nm, Data interval: 1.0 nm, Integration time (step): 0.5 s

E. Each sample was scanned three consecutive times. The raw CD spectra were processed using Pro-Data Viewer software. Final spectra for tubes 3 and 5 were baseline-corrected by subtracting the spectrum of tube 1 (blank), and spectra for tubes 4 and 6 were corrected by subtracting the spectrum of tube 2 (STX alone).

### 4.9. Safety and biological activity of circular aptamers assessed in mice

The biological activity of the circular aptamer was evaluated via tail vein injection using a mouse biological assay described by the Association of Official Analytical Chemists (AOAC) in the USA in 2005. ICR mice (4 weeks old, 18-20 g, half tail vein injection for each mice) were used to observe survival rates. Based on previous experience, if the mice did not show any abnormalities within 10 min of STX injection, they were not expected to die, and their clinical signs would not differ from those of normal mice within 48 h of observation. Therefore, mice that did not die within 10 min of STX injection were considered survivors.

For tail vein injection, the veins on the left and right sides of the back of the tail were chosen.

#### 4.9.1. Exploration of lethal dose in mice

Based on a previous study, four concentration gradients of STX (12 μg/kg, 11.25 μg/kg, 10.5 μg/kg, 9.75 μg/kg) were used to explore the lethal dose of STX in mice. Six mice in each group were observed for mortality, and an STX dose near LD_100_ was selected for further experiments.

#### 4.9.2. Evaluation of the biological activity of circular aptamers

Preliminary acute toxicity tests and pharmacodynamic evaluations were also performed. The molecular weight of the Aptamer-STX complex (7.6–8.8 kD) is significantly greater than that of the STX alone (299 Da). Therefore, the 3-kD ultrafiltration membrane can theoretically retain the aptamer-STX complex, and only the unbound STX would pass through the membrane and be free in the effluent, as shown in Table 6, where the concentration of STX in experimental groups 5-10 is 11.25 μg/kg. Therefore, in this study, STX solution was prepared at an international dose of 10 mL/kg, mixed with the aptamer at a 1:1 ratio with regard to molecular number, and incubated at room temperature(around 27℃) for 1 h.

After incubation, the mixture was injected into the tail vein of 6 mice in each group.

**Table 2.** Grouping design of caudal venous injection experiment in mice

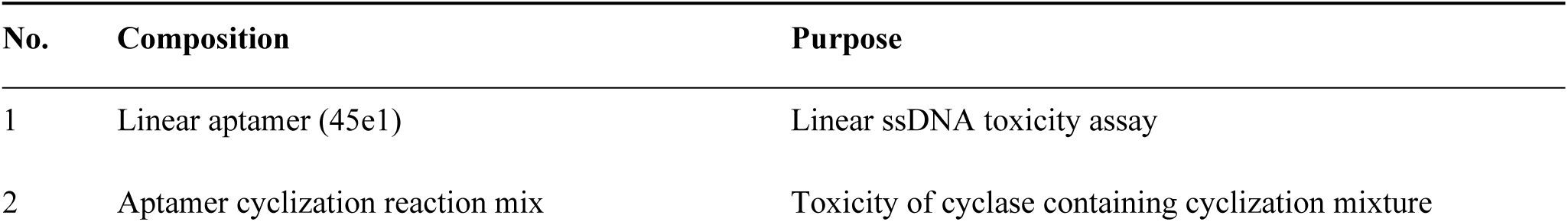
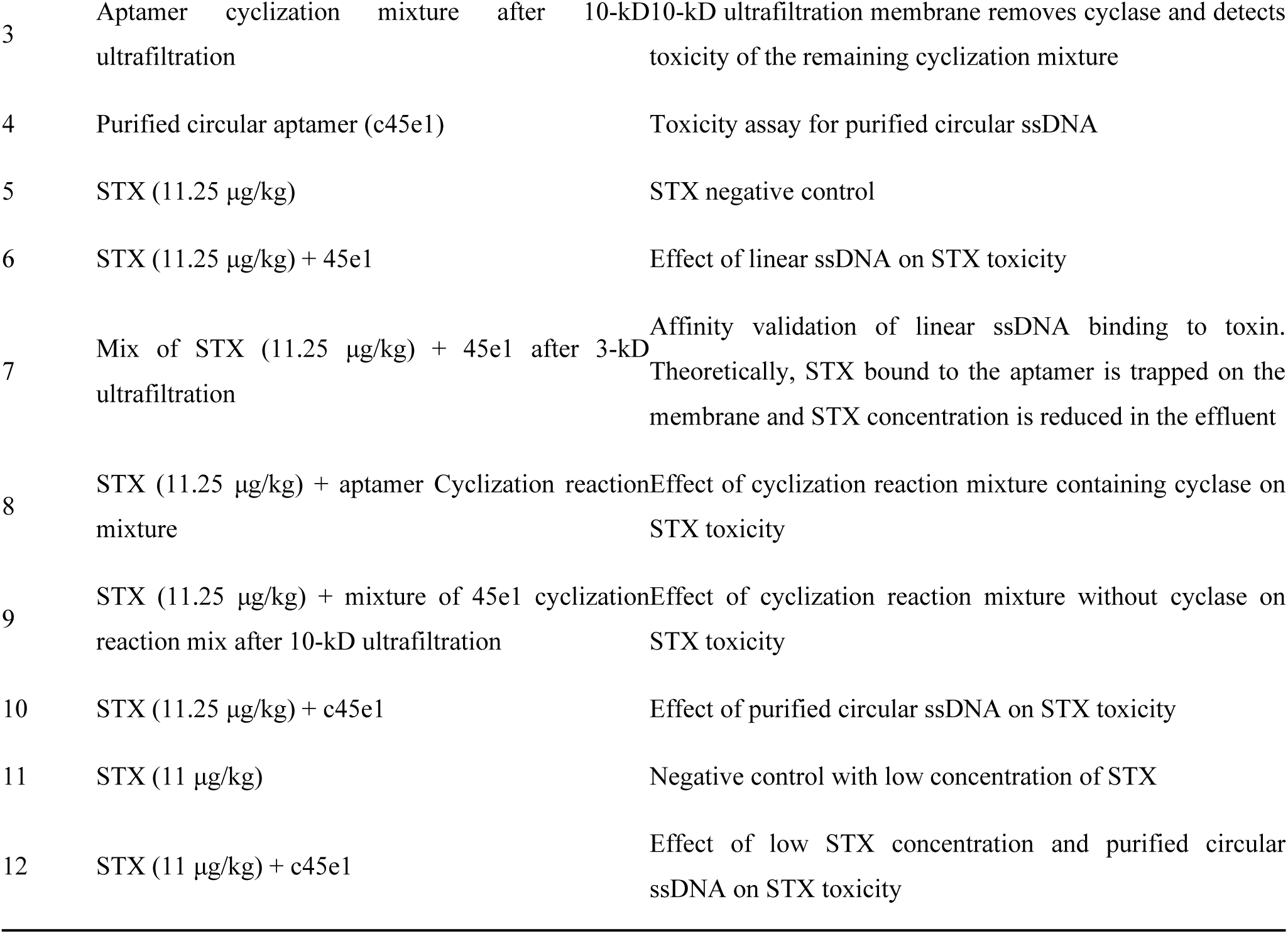

#### 4.9.3 Statistical analysis

The data were analyzed using one-way ANOVA to determine the significance of the differences between the mice groups, p < 0.01 was consider as significant result.

## Acknowledgments

This work was supported by the National Natural Science Foundation of China [grant number 82173732] and National Key R&D Program of China [grant numbers 2019YFC0312600 and 2019YFC0312603].

## Author contribution

Conceptualization, L.W.; M.S.; H.C; methodology, S.G. and B.D.; software, S.G, B.D and D.C; Y.W; validation, S.G., B.X. and Y.J.; formal analysis, H.Y. and B.D.; investigation, T.L., G.L., Y.Z., C.J.; resources, L.W.; data curation, S.G.; writing—original draft preparation, S.G.; writing—review and editing, S.G., B.D. and D.C.; visualization, B.D.; supervision, H.C., Z.M., Y.G., H.Z.; project administration, M.S.; funding acquisition, L.W. All authors have read and agreed to the published version of the manuscript

