## Supporting Information for "Computational Assessment of Circularization Impact on Saxitoxin G-quadruplex Aptamer Performance"

*1. Modeling the 3D structure of 45e1 and c45e1*

1.1. 3D Structure of 45e1

The sequence with the highest score, (5′-GGGCGCGGTGATCGGAGAGGG-3′), is chosen to build the G-quadruplex core model with 3D-Nus. Two G-quarters were formed from sequences G6, G12, G20, G26, G7, G13, G21, and G27, as shown in Fig. S1a. Nine possible structural models, including parallel, antiparallel, and mixed types of the G-quadruplex, were modeled using 3D-Nus, s shown in Fig. S1b. Molecular dynamics at a constant temperature of 300 K for 10 ns after the addition of $Mg^{2+}$ to the center of the G-quadruplex using showed that the (Root mean square deviation)RMSD values for Q3 (parallel type) were consistently lower than those of the remaining eight conformations, demonstrating that Q3 is more stable than the other nine structures and most likely the natural structure of 45e1. The variation in RMSD is shown in Fig. S1c. Therefore, Q3 was used as the 3D structure of 45e1 for subsequent studies (see Fig. S1d).

**Table S1.** The most stable quadruplex sequence of 45e1 predicted by QGRS Mapper

| position | Length | QGRS (5′—3′) | G-score |
|---|---|---|---|
| 6 | 22 | **GG**GCGC**GG**TTGATC**GG**AGAG**GG** | 19 |

The bolded bases are those involved in the formation of the quadruplex

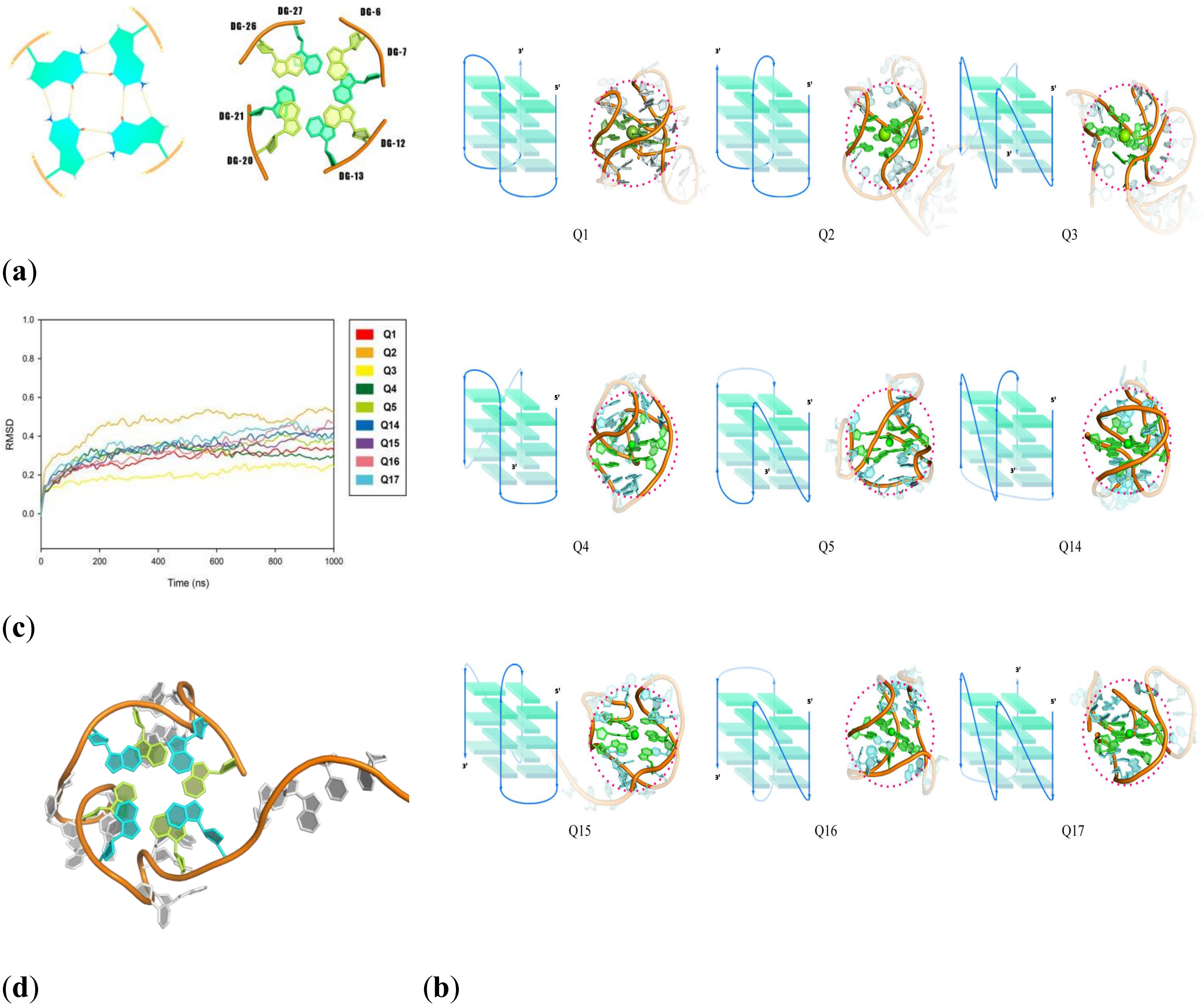


**Fig. S1.** Process of modeling the 45e1 3D-structure. (a) Two-layer G-planes formed by the G-core sequences portion of 45e1 (b) Nine possible 3D structures of 45e1 (containing Q1, Q2, Q14, and Q18 as antiparallel structures, Q3 as a parallel structure, and Q4, Q5, Q15, Q16, and Q17 as mixed structures) (c) Variation in the RMSD curves of molecular dynamics for the nine 3D structures at a constant temperature of 300 K for 10 ns (d) 3D structure of 45e1 in Q3 (the four guanines in blue and green represent guanines that make up the different G-planes)

1.2. 3D structure of c45e1

Circular 45e1 modeled using OxDNA was visualized via PyMOL, and the results are shown in Fig. 2. The original G-planes were disrupted and the G-core sequences were spatially distant, making it difficult to form a G-quadruplex structure. This confirmed our previous finding that cyclization may disrupt the original 3D structure of the aptamer. Whether the current structure can maintain its original high affinity requires further verification using molecular docking and dynamics.

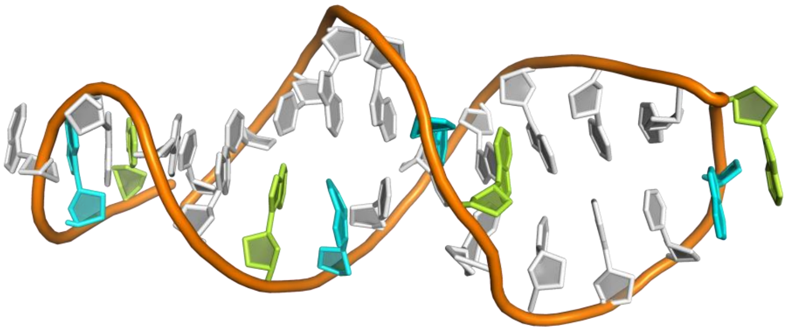

**Fig. S2.** 3D structure of c45e1. The blue and green bases are components of the two G-quarters in 45e1

*2. Cyclization of linear 45e1*

2.1. Small-scale cyclization of linear 45e1

Due to the conformational changes in c45e1, they are subjected to different migration resistances during electrophoresis compared to 45e1 and therefore, show different apparent molecular weights on PAGE (Fig. S2). There was an obvious difference in the migration rates between cyclized and uncyclized 45e1. It is evident from lanes 1 and 2 that plain 45e1 was degraded by Exo I, whereas lanes 3 and 4 show that the band representing c45e1, located below that of 45e1, was resistant to degradation by Exo I. It can be concluded that c45e1 has a faster migration rate owing to its lower resistance to electrophoresis after cyclization, and that its electrophoretic band is located more anteriorly compared to the 20-nt position.

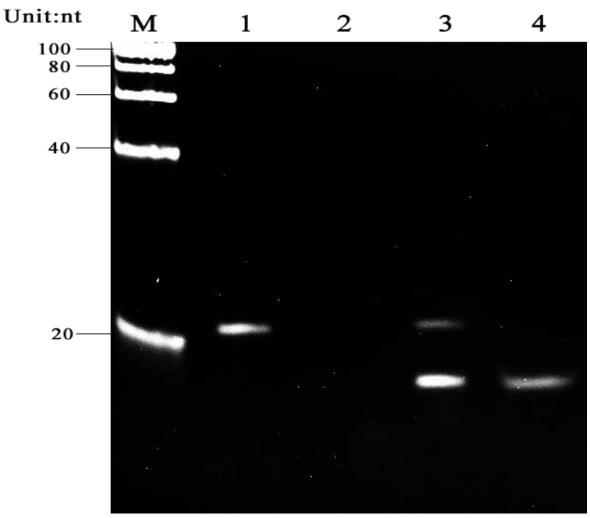


**Fig. S2.** Results of cyclization of linear 45e1. M: Marker; 1: ssDNA (27 nt); 2: ssDNA + Exo I; 3: ssDNA + CircLigase (100 U); 4: ssDNA + CircLigase (100 U) + Exo I.

2.2. Bulk preparation of c45e1

For the bulk preparation of c45e1, substrate was added in excess and 1.5 mm of 15% PAGE electrophoresis was used. Results are shown in Fig. 8. The brightness of the uncyclized band increased when the substrate was amplified from 60 μL to 100 μL, indicating that the reaction system was expanded, and the efficiency of cyclization reduced. Simultaneously, another band appeared more clearly between 40 and 60 nt. Because of the large apparent molecular weight, it is speculated that this may be due to an increase in substrate density during the bulk cyclization reaction, with some of the substrates interacting or becoming entangled, or due to the formation of multi-loop-linked or tandem polysupercircular DNA during DNAligase activity. c45e1 was only considered a monomer in this study; therefore, only the band corresponding to 20 bp was collected for subsequent validation, and the monomeric c45e1 was purified for Bio-layer interferometry(BLI) experiments.

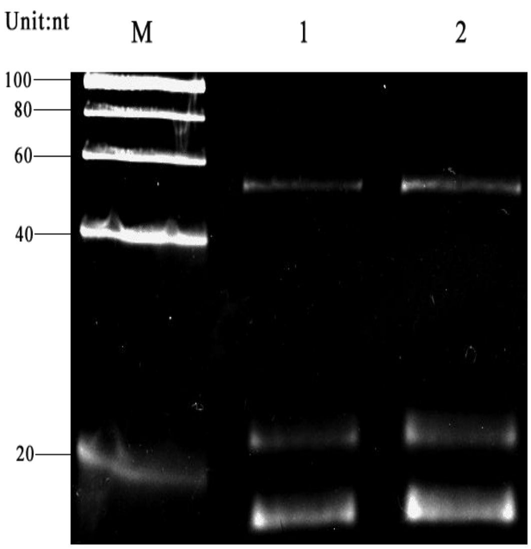


**Fig. S3.** Results of bulk preparation of c45e1. M: Marker; 1: 60 μL of 45e1 (27 nt) added; 2: 100 μL of 45e1 (27 nt) added.

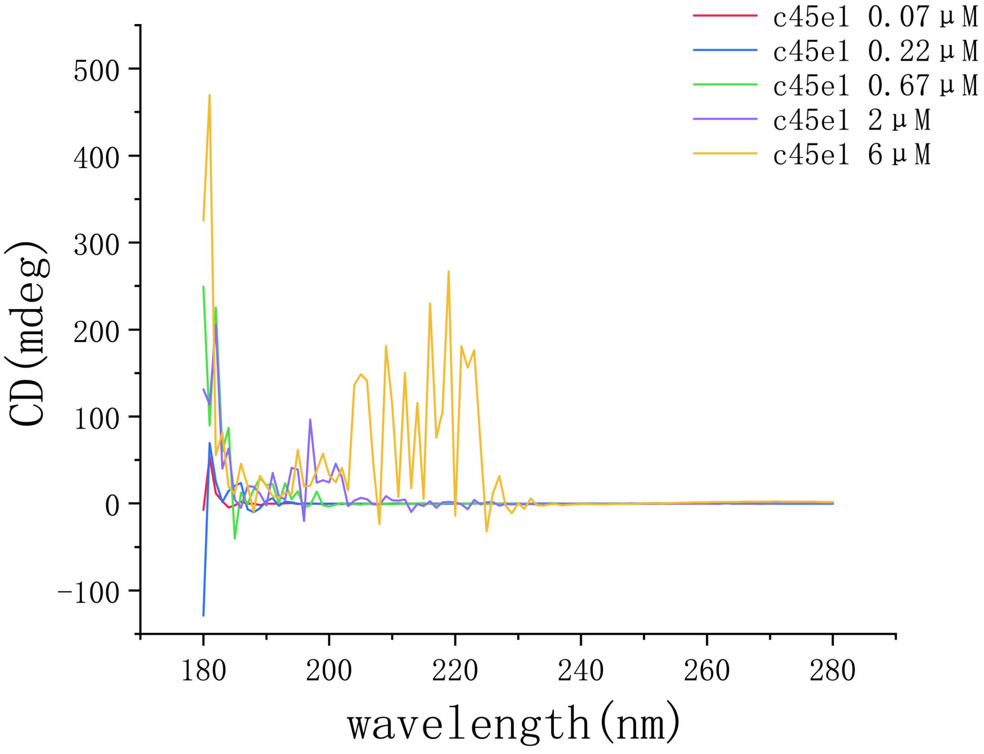


**Fig. S4.** Different concentration of c45e1 in CD spectrum.
